# Analysis of ipsilateral corticocortical connectivity in the mouse brain

**DOI:** 10.1101/241125

**Authors:** Akiya Watakabe, Junya Hirokawa

## Abstract

In primates, proximal cortical areas are interconnected via within-cortex “intrinsic” pathway, whereas distant areas are connected via “extrinsic” white matter pathway. It is not well known how cortical areas are interconnected in small-brained mammals like rodents. In this study, we systematically analyzed the data of Allen Mouse Brain Connectivity Atlas to answer this question and found that the ipsilateral cortical connections in mice are almost exclusively contained within the grey matter with the exception of the retrosplenial area. We analyzed the layer-specific distribution of axonal projections within the grey matter using Cortical Box method and obtained the following results. First, widespread axonal projections were observed in both upper and lower layers in the vicinity of injections, whereas highly specific “point-to-point” projections were observed toward remote areas. Second, such long-range projections were predominantly aligned in the anteromedial-posterolateral direction. Third, in majority of these projections, the connecting axons traveled through layer 6. Finally, the projections from the primary and higher order areas to distant targets preferentially terminated in the middle and superficial layers, respectively, suggesting hierarchical connections similar to those of primates. Overall, our study suggests the conserved nature of neocortical organization across species despite conspicuous differences in wiring strategy.

## I. INTRODUCTION

Expansion of the cerebral cortex and its differentiation into many specialized areas is one of the most distinguished features of the mammalian brain evolution. The connections among these cortical areas lay the basis for functional integrity of the cortex as a whole. In macaques and other non-human primates, anterograde tracer experiments found two modes of cortical connections: the distant areas are connected via bundles of axons that travel through the white matter (“extrinsic” connection), whereas the nearby areas are connected via “intrinsic” or “horizontal” cortical connections that are contained entirely within the grey matter (Levitt et al. 1993; Lund et al. 1993). How, then, are cortical areas connected in small-brained animals like mice? Although much smaller in size, the mouse cortex also has complex cortical organization (Horvá et al. 2016; Goulas et al. 2017). The visual system, for example, has multiple higher visual areas that each has unique inter-areal connectivity (Wang and Burkhalter 2007; Wang et al. 2012). It is possible that they are interconnected via the white matter as in primates. On the other hand, the whole mouse cortex is much smaller than macaque V1 (Finlay 2016). In terms of size, the mouse cortex may exhibit intrinsic-type connectivity.

As expected from its importance as a model animal for neuroscience studies, there have been many efforts to elucidate the cortical connectivities in rodents (e.g., Schü et al. 2006; Oh et al. 2014; Paxinos 2014; Zingg et al. 2014; Horvá et al. 2016). However, there has been little mention in the literature on where axonal fibers actually pass through. In a classic single cell tracking study, Dechenes and co-workers showed several examples of rat S1 neurons projecting to S2 via the grey matter (Zhang and Deschêes 1997). Likewise, we labeled the callosally projecting M1 neurons by double viral strategy and found their collateral projections to reach the ipsilateral S1 via the grey matter (Watakabe et al. 2014). These observations raise the possibility of grey matter pathway for mouse cortical connections but they are too fragmental to generalize: other cell types may project to distant targets via the white matter even in S1 or M1 and/or there may exist white matter connectivity between other areas. To clarify this point, there is a need for systematic analyses of the route of axonal projections in rodents.

In this article, we approached this question by using a publicly available database of Allen Institute for Brain Science (http://connectivity.brain-map.org/). In this database, there exist serial image data of hundreds of samples, in which AAV-based anterograde tracers were injected at a single site per mouse. Examination of the fiber tracts in their dataset revealed that most of the ipsilateral cortico-cortical connections are explained by spread of axon fibers within the grey matter. We also quantitated layer-specific distribution of the axonal signals by modified Cortical Box method, which is a kind of a flat map with laminar information (Hirokawa et al. 2008a; Hirokawa et al. 2008b; Watakabe 2012). Combined with clustering analysis, it provided an objective means to investigate the complex architecture of axonal projections within the grey matter. Based on these analyses, we clarified common aspects of cortical wiring across species, despite differential segregation patterns to grey and white matter.

## II. MATERIALS AND METHODS

### Confocal microscopic analyses

For confocal microscopic imaging of the cortico-cortical fibers, we used the brain sections that we used in our previous study (Watakabe et al. 2014). Specifically, we used the section of mouse #456, in which NeuRet vector encoding TRE-SYP_CFP was injected into S1 and AAV encoding Syn-rtTAV16 was injected into M1. The sections were pretreated with 80 % methanol/20 % dimethyl sulfoxide solution (Dent’s solution) and immunostained with the anti-GFP antibody (1:20000, Tamamaki et al. 2000) followed by Cy2 conjugated anti-rabbit IgG 1:1000, as described previously (Watakabe et al. 2014). After mounting on a slideglass, the stained sections were imaged by Olympus Fluoview FV1000 confocal microscopy using 40X water immersion lens.

### Digital data acquisitions

We downloaded a series of dataset from a public database of mouse connectome at www.alleninstitute.org (Allen Mouse Brain Connectivity Atlas (2011)). They conducted a brain-wide anterograde tracing with enhanced green fluorescent protein (EGFP)-expressing AAV in mice (Oh et al. 2014). Seventeen datasets shown in Table 1 were selected for detailed analyses, which include areas with various functions and topological positions. We used only the wild type mice, because we wanted to examine all the potential projections from the area of interest. The XY resolution of the original dataset was 0.35µm/pixel and the imaging was performed with 100 µm intervals in Z direction. For high magnification views, we used the original resolution images. For other purposes, including Cortical Box analysis, we downloaded 1/8 compressed version of 8bit RGB, JPEG format images without changing the default contrast for each image.

**Table 1.**
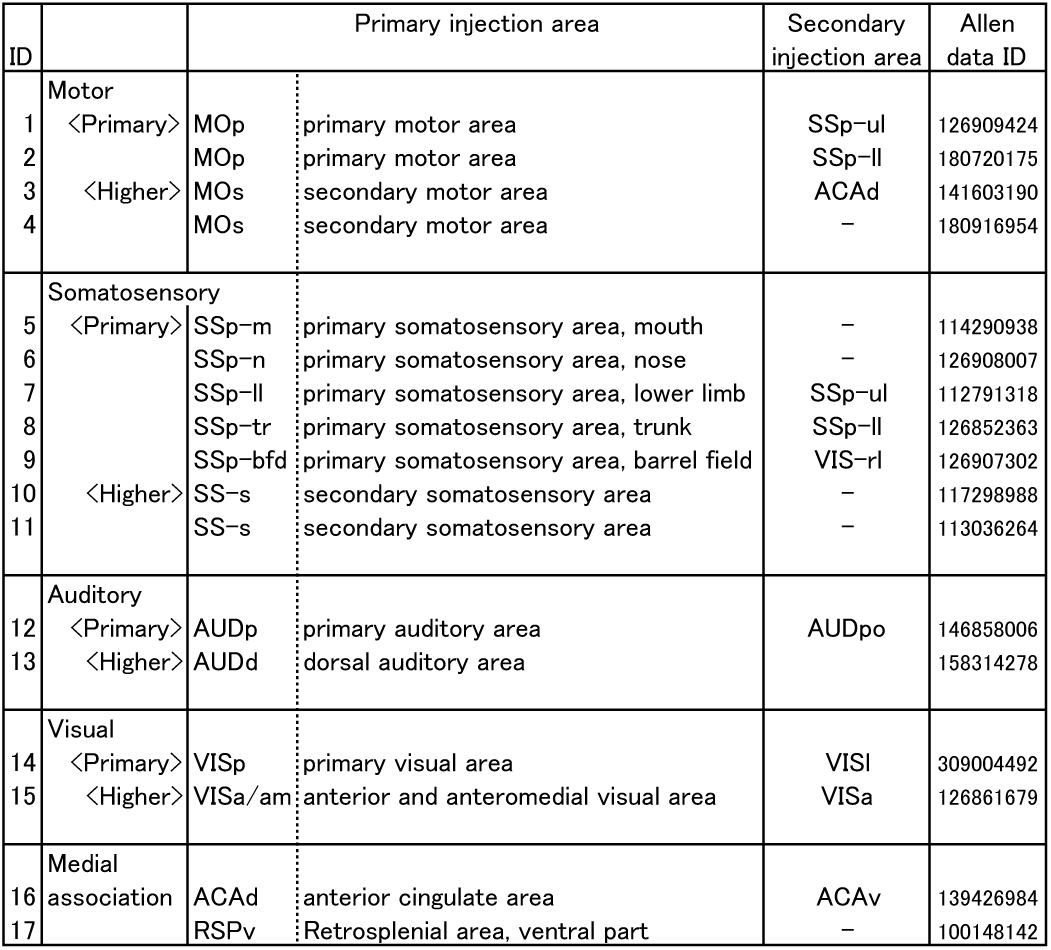
Analyzed areas in the present study.

### Standardization of regions of cortex

The cortical box method was performed as previously described (Hirokawa et al. 2008a; Hirokawa et al. 2008b; Watakabe 2012). The cortical regions defined by the medial and lateral ends were cut out from the images and transformed to rectangles. The medial end of the white matter and the valley of the rhinal fissure were chosen as structural landmarks of the mediodorsal (MD) and lateroventral (LV) ends of the cortical sections, respectively. The border between the cortex and the white matter were chosen as the grey matter-white matter boundary. The pial surface and the bottom of the white matter area was used as the outer (OC) and the inner contours (IC), respectively. In Fig. 7, we extended the analysis to the area around the medial wall. The bottom of the medial wall and the symmetric counterpart in the cortex were manually chosen as the medial (ME) and lateral (LE) ends. Note that we only used the medial box method to visualize individual projections without comparing across samples, since the method does not allow objective definition of the ROI. To reduce the effort to make cortical box from sections without signals, we used 21–57 coronal sections in each projection so that we can cover the injection site and the major projections in each projection.

To allow comparison of the cortical boxes between different samples, we aligned the anterior-posterior positions based on the shape of the hippocampus in reference to the Paxinos and Watson (Paxinos and Franklin 2004). We limit the analysis in the range between Bregma distance of +1.5 to -4.6, spanning from primary visual cortex to motor cortex with a resolution of 100 μm. Therefore, a standard cortical box consists of three dimensional data; 61 × 1000 × (100+30) data points (AP length × ML width × (layer depth and white matter fraction) with the relative intensities of fluorescent signals expressed with percentages (0–100%). To generate the standardized map of a particular layer, the specific layer fraction (1–10% for layer 1; 75–100% for layer 6) and white matter fraction were extracted from the standardized cortical box and compressed into a two-dimensional map by averaging the fraction with a Gaussian filtering using a kernel of 7 pixels. Visualizations were carried out using Matlab 2017 (Mathworks, Natick, MA, USA).

### Annotation of cortical areas

To annotate the cortical areas, we entirely relied on the annotation by the Allen Mouse Brain Connectivity atlas. This database provides two ways to annotate cortical areas. First, in their “cortical map signal viewer”, they provide flatmap version of the cortical signals with annotation. We were able to identify the “hotspot” with concentrated tracer signals by comparing their map with coronal images and with our Cortical Box views. Second, their atlas is coupled with the Nissl-based reference atlas. We annotated the cortical areas based on such information. Thus, the nomenclature of the areas in our study is in accordance with that of the Allen atlas.

### Data analysis of standardized cortical map

For clustering analysis, we compressed the cortical box by downsampling the ML width from 1000 points to 100 point with nearest neighbor method to reduce the computational burden. Columns with low signals (mean intensity less than 50) were removed from analysis. The rest of the columns from 17 cortical boxes were pooled and the standardized layer maps were analyzed as a P x N matrix (row × column), where P is the location number (P=24,000 points), and N is the number of cortical layer data (N=100). Each column of this matrix was normalized using the maximum and minimum of the data in that column. The data matrix was subjected to hierarchical cluster analysis using Ward's algorithm. The number of the cluster was systematically changed and the variance from the average was calculated for each cluster and summated to calculate “total within sum of square”, which provided the measure of similarities within the cluster (Fig. 9A). Based on this evaluation, we set the cluster number to five and performed the subsequent analyses.

To test the robustness of the result of clustering, we performed clustering using a subset of the 17 injections and calculated the similarity of thus-obtained patterns with the original patterns. In more detail, we randomly chose a subset (N=1–16) of the 17 samples and performed the clustering analysis with a fixed cluster number (n=5). Each clustering analysis generated five averaged layer patterns and they were compared with the original five patterns (Fig.9B) for the best matching. Then, the Pearson's correlation coefficient was calculated between the new and the original patterns and five of them were averaged as the similarity value of the new clustering as a whole. We repeated 20 times of random sampling for each sample number (N=1–16) and the average of the 20 similarity values were plotted against the sample number (shown in red in Supplementary Fig.3). For comparison, we also calculated the correlation coefficients among the five patterns generated for each clustering and the highest values were averaged for each sample number (shown in blue in Supplementary Fig.3). Supplementary Fig.3, thus, also indicates that the similarity across the clusters is low, even if the sample numbers are reduced.

To visualize the spatial distributions of thus-classified columnar lamina patterns, each point on the flatmap was color-coded for five patterns and remapped on the layer map for each injection. Overlaid on this color-coded map was the maximum intensity projection of all layers (shown as contours in Supplementary Fig.2). This MIP map was used to manually track the local maximums from the injection site to the two furthest target areas. For cross-sample analyses, only these two axon pathways were extracted (Fig. 9D) for remapping on a common coordinate (see above) (Fig. 9E).

To evaluate the difference of the target layer patterns for the projections from the primary and higher order areas, we calculated the feedforward index (b/(a+d)), which compare the middle-layer dominant projections to superficial layer dominant projections. The significance of the difference of the feedforward indices between the primary and the higher order areas were validated by bootstrap method (10000 repetitions) by re-assigning primary and higher order areas pseudo-randomly.

### Data and statistical analyses

The primary data of Serial Two-photon tomography was obtained from Allen Mouse Connectivity atlas (www.alleninstitute.org). The datasets generated during and/or analyzed during the current study are available from the corresponding author on reasonable request. The statistical significance of the bias in the direction for long-range projections was determined by Kuiper test. The statistical significance of differential pattern distribution for the primary and higher order areas were tested by bootstrap method (Table 2; see above). Student's t tests were performed for comparison of the pattern distribution for the primary and higher order areas in Supplementary Fig.3A. Matlab was used for calculation and all values are expressed as means ± SEM.

**Table 2.**
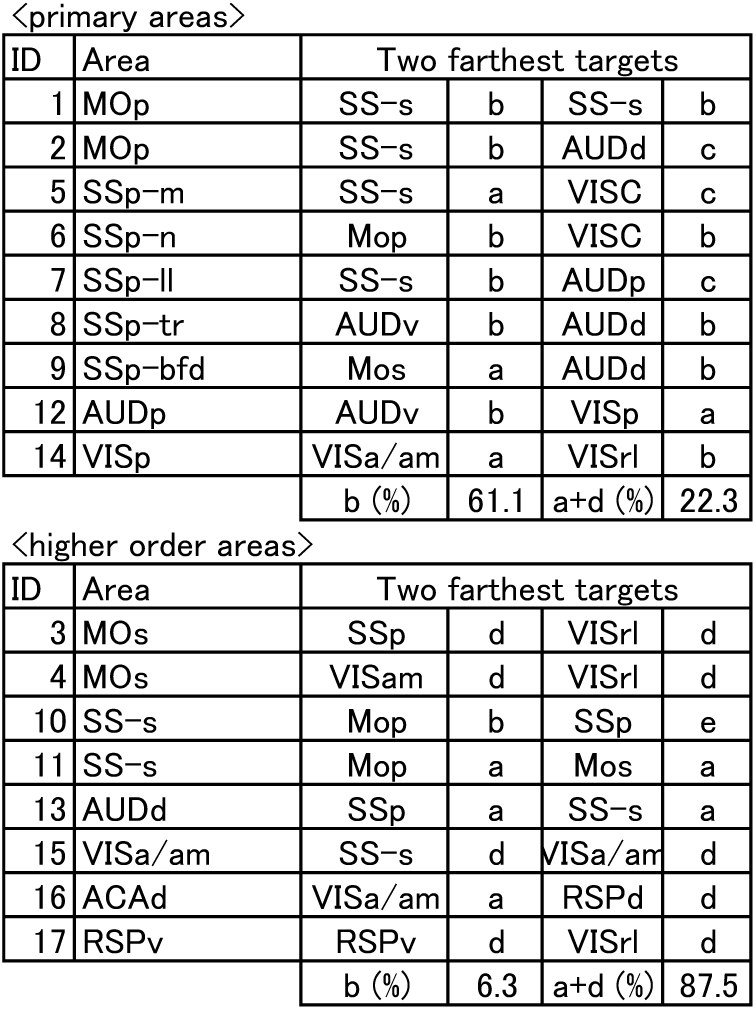
Laminar preference of innervation at remote targets.

| <primary areas> |  |  |  |  |  |
| --- | --- | --- | --- | --- | --- |
| ID | Area | Two farthest targets |  |  |  |
| 1 | MOp | SS-s | b | SS-s | b |
| 2 | MOp | SS-s | b | AUDd | c |
| 5 | SSp-m | SS-s | a | VISC | c |
| 6 | SSp-n | Mop | b | VISC | b |
| 7 | SSp-ll | SS-s | b | AUDp | c |
| 8 | SSp-tr | AUDv | b | AUDd | b |
| 9 | SSp-bfd | Mos | a | AUDd | b |
| 12 | AUDp | AUDv | b | VISp | a |
| 14 | VISp | VISa/am | a | VISrl | b |
|  |  | b (%) | 61.1 | a+d (%) | 22.3 |
| <higher order areas> |  |  |  |  |  |
| ID | Area | Two farthest targets |  |  |  |
| 3 | MOs | SSp | d | VISrl | d |
| 4 | MOs | VISam | d | VISrl | d |
| 10 | SS-s | Mop | b | SSp | e |
| 11 | SS-s | Mop | a | Mos | a |
| 13 | AUDd | SSp | a | SS-s | a |
| 15 | VISa/am | SS-s | d | VISa/am | d |
| 16 | ACAd | VISa/am | a | RSPd | d |
| 17 | RSPv | RSPv | d | VISrl | d |
|  |  | b (%) | 6.3 | a+d (%) | 87.5 |

## III. RESULTS

### Axonal fibers that project from M1 to the ipsilateral S1 are contained within the grey matter

In our previous double viral experiments, we examined the ipsilaterally projecting axon collaterals of callosaly-projecting M1 neurons (Watakabe et al. 2014). To generalize our finding, we first examined the axonal fibers between M1 and S1 of the ipsilaterally-projecting M1 neurons. For this purpose, retrograde lentiviral vectors harboring fluorescent marker protein (SYP-CFP) under the TRE promoter was injected in S1, whereas TET activator in AAV was injected in M1 (Fig. 1A). Ideally, the labeled axon fibers should represent all kinds of neurons that link M1 and S1 (Fig. 1B-G). In this labeling, the axonal fibers that connect M1 and S1 were entirely contained within the grey matter (Fig. 1C). This observation suggests that all the direct connections from M1 to S1 are through axons within the grey matter. At the terminal region, axon fibers branched into numerous small fibers and formed bouton-like varicosities at S1 (Fig. 1C, F and G). In contrast, the axon fibers that proceed within layer 6 were generally thicker and had few axon branches, if any (Fig. 1D and E). Nevertheless, we could also observe fine fibers with varicosities to coexist with such thick fibers, suggesting the existence of synaptic connections during the layer 6 pathway.

**Fig. 1.**
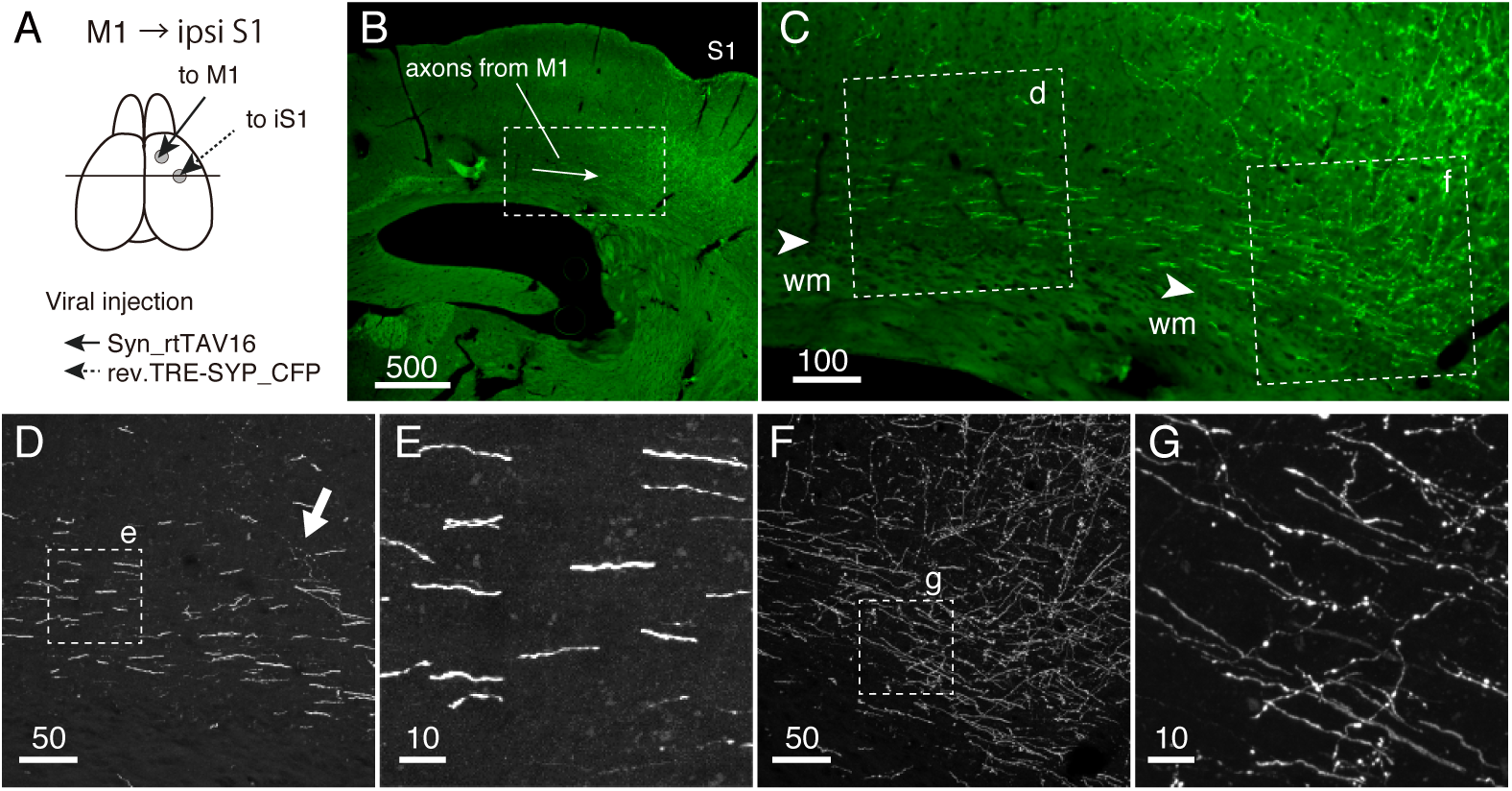
Confocal imaging of the cortico-cortical axon fibers. The cortico-cortical axons that project from the motor cortex (M1) to the ipsilateral somatosensory cortex (ipsiS1) were selectively labeled for confocal imaging. (A) Schematic representation of the labeling strategy. Simultaneous injections of the NeuRet vector coding TRE-SYPCFP into ipsiS1 and AAV coding Syn_rtTAV16 resulted in labeling of all types of connections between the two areas. (B) Low magnification view of the section immunostained by GFP antibody showing labeled axons from M1 (shown by an arrow). The rectangle region is magnified in panel C. Bar: 500 μm. (C) Higher magnification view of M1-to-S1 axons. Note that the border between Layer 6 and the white matter (wm) can be unambiguously identified (arrowheads). Two regions shown by dotted squares with labels d and f were imaged by the confocal microscopy in panels D and F. Bar: 100 μm. (D) Maximum intensity projection image of thick axon fibers that reside at the very bottom of layer 6 in panel C. The white arrow indicates the intermingling of thin axon fibers with bouton-like varicosities. Bar: 50 μm. (E) Magnified view of panel D. Note the lack of side branches or boutons. Bar: 10 μm. (F) Maximum intensity projection image of terminal arborization at ipsiS 1. Note numerous fine branches to come out of layer 6 toward upper layers. Bar: 50 μm. (G) Magnified view of panel F. Note the presence of bouton-like varicosities. Bar: 10 μm.

### Examination of AAV-based anterograde tracers of Allen Mouse Brain Connectivity atlas

To elaborate our finding, we made use of Allen Mouse Brain Connectivity atlas (http://connectivity.brain-map.org/). Among the dataset, we selected only those using the wild type mice and AAV with synapsin-promoter driven EGFP. This AAV tracer works as an efficient anterograde tracer and labels cell bodies, dendrites, axonal fibers and terminals efficiently. The sections are imaged by serial two-photon tomography with 100µm interval (Ragan et al. 2012; Oh et al. 2014). We browsed through the dataset and selected 17 injections for detailed analyses (Table 1). These injections were selected based on (1) high level of labeling, (2) high quality of imaging and (3) variety of cortical areas to represent both primary and higher order areas from the motor, somatosensory, auditory and visual systems. Although this AAV tracer labels all the cortical projection neurons unlike the double viral system that we employed for Fig. 1, we reasoned that we should be able to distinguish cortico-cortical projections from other projections by tracking the fibers across serial sections.

Fig. 2 shows an example of injection into the dorsal auditory area (AUDd). In this example, there was a robust projection to the somatosensory cortex (Fig. 2E, SSp). At first sight, it may appear that thick axon fibers in the white matter convey this connection. At higher magnification, however, the axon fibers in the white matter rarely branched into the grey matter within the ipsilateral hemisphere (Fig. 2C and G). Instead, they proceeded through the corpus callossum without losing fluorescent intensity toward the contralateral side (Fig. 2A and E, arrowheads) where they branched into thinner fibers to innervate the contralateral cortices (Fig. 2A; asterisks, Fig. 2I). Based on comparison of the fluorescence intensity and thickness of the axon fibers that just branched out from the injection site (Fig. 2A, arrow) and those that reached the contralateral hemisphere (Fig. 2A and E, arrowheads), it appears that the majority of the axon fibers in the white matter innervate the contralateral side. Where, then, do the ipsilateral cortical projections pass through?

**Fig. 2.**
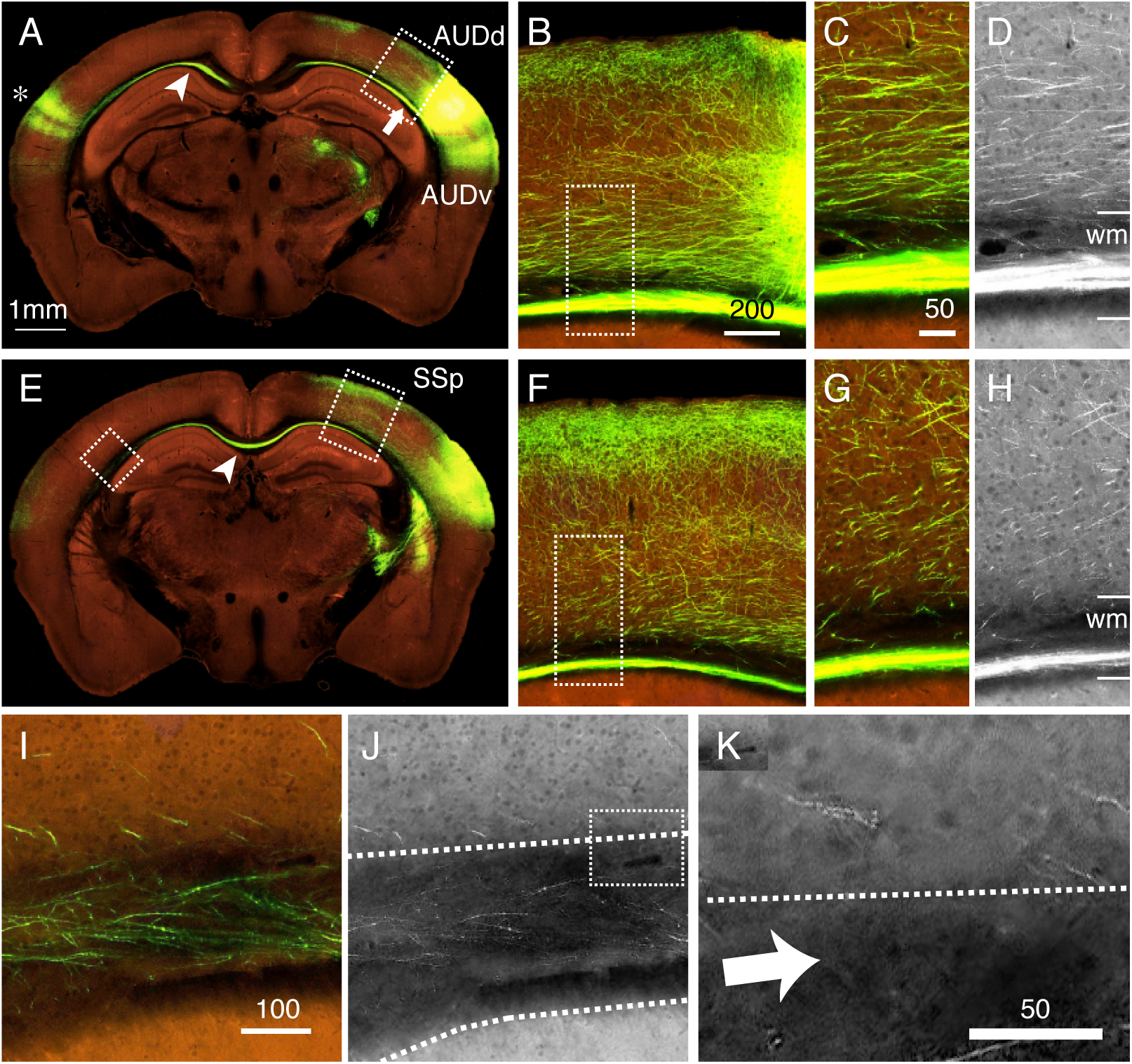
Image data from Allen Mouse Brain Connectivity atlas analyzed for the axonal projections. The AAV tracer was injected into AUDd (Table 1: ID13) and the axonal spread of GFP signals were imaged by Serial two-photon tomography. (A) The coronal plane with center of injection. The white arrow indicates the axon bundle within the white matter that projects towards the contralateral area (shown by asterisk). Note that the intensity of the fluorescence does not change much, when compared between the intensity right after entering the white matter (arrow) and after entering the contralateral hemisphere (arrowhead). The dotted box is magnified in panel B. Bar: 1 mm. (B) The magnified view of axons projecting from the injection site. Note that, few, if any, axon fibers exist between the white matter bundle and the fibers in the bottom of layer 6. The dotted box is magnified in panel C. Bar: 200 μm. (C) Magnified view around the border between layer 6 and the white matter. Bar: 50 μm. (D) Red channel view of the image shown in panel C. This channel shows the autofluorescence background of the tissues with some leak of green tracer signals. The border between the white matter (wm) and layer 6 was determined by darkness of white matter and the cell shadows (see methods). (E) The coronal section view, which is anterior to that in panel A. The arrowhead indicates the axon fibers crossing the corpus callosum. Note the axonal projections to upper layers in SSp. Magnification is the same as in panel A. (F) Magnified view of the dotted box around SSp of panel E. Magnification is the same as in panel B. (G) Magnified view of the dotted box of panel F. Magnification is the same as in panel C. (H) Red channel view of the image shown in panel G. (I) Magnified view of the dotted box in the contralateral hemisphere of panel E. Note the presence of axon fibers within the white matter. Bar: 100 μm. (J) Red channel view of the image shown in panel I. (K) Magnified view of the dotted box in panel J. The dotted line shows the border between layer 6 and the white matter. Note the presence of cell shadows in layer 6. The border is determined by the overall autofluoresence intensity. So, the presumptive white matter region may include some “layer 6”, because we can see very weak cell shadows (white arrow). In order not to underestimate the “white matter”, we prioritized the overall fluorescence intensity over cell shadows in border determination.

In this case of AUDd injection, ventrolateral projections to AUDv and anteromedial projections to SSp were present (Fig. 2A and 2E). In the former case, AUDv, which lies next to AUDd, received side-by-side projections within the grey matter and no axons went through the white matter (Fig. 2A). Similarly, numerous axon fibers projected out medially in the vicinity of the injection site, which were directed parallel or oblique to the cortical layers (Fig. 2B). In particular, relatively thick fibers populated within layer 6, which traveled across sections to reach the bottom part of SSp, where they branched into thinner fibers, turned perpendicular to the layers and innervated layers 1–3 (Fig. 2F). There were also fibers that proceeded through other layers to reach SSp. These observations again showed that ipsilateral cortical connections are established through widespread axon fibers that are contained within the grey matter and not through the white matter.

One key point of observation for the above conclusion is the distinction between the white matter and the grey matter. In this and the subsequent studies, we drew borders based on the intensity of the tissue autofluorescence in the red channel (Fig. 2I-K). With this criterion, the “bright” region is definitely a grey matter, because the shadows of cell bodies observed at high magnification tell us that the region is densely populated with neuronal cells. The “dark” region may include some “grey matter”, but we classify such region as “white matter” in order not to underestimate the white matter region.

### “Grey matter route” is a rule rather than an exception for cortico-cortical connections

To elaborate the observations obtained from AUDd injection, we examined other cortical projections. Fig. 3 illustrates twelve typical cases of cortical projection patterns (Cyan arrows indicate the overall direction of the axon tracts). Fig. 3A-F exhibit the cases in which axons projected toward the areas on the lateral side. In all these cases, we observed axon fibers that travel through layer 6 toward the target area, where they turned oblique to the laminar planes and terminated in a columnar fashion. The thick axon bundles that entered the white matter either turned medially to project to the contralateral cortex or entered into the internal capsule. In none of these cases, thick axon bundles extended toward the target cortical area. In some instances, we observed relatively thin axon fibers to go through the white matter ventrolaterally (e.g., Fig. 3F). In such cases, axon fibers appeared to enter the striatum but not the cortical grey matter (e.g., Fig. 4E, lower arrow).

**Fig. 3.**
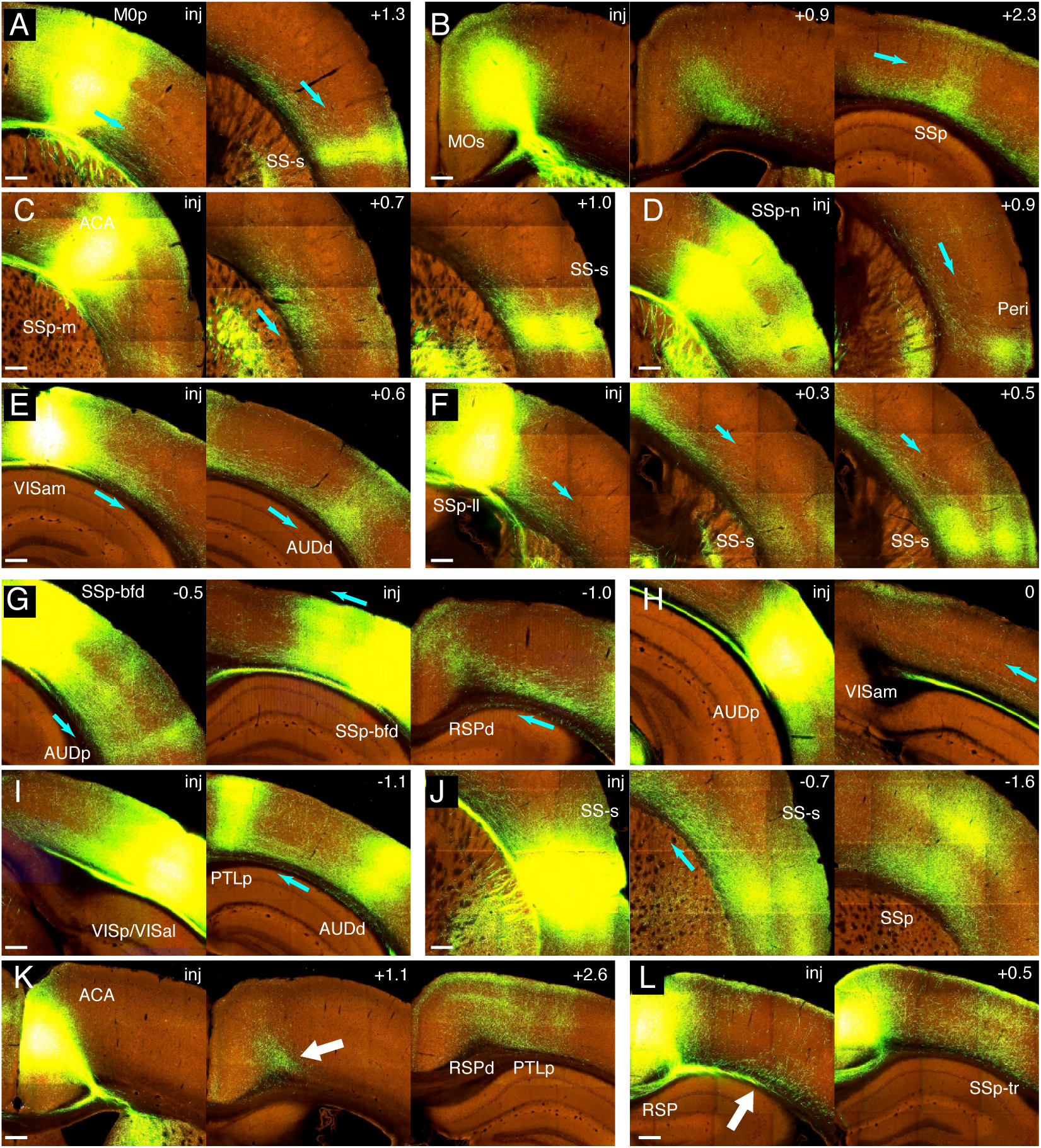
Example images of Allen Mouse Connectivity atlas dataset showing “grey matter route” of various cortico-cortical projections. Panels A-L shows a set of images showing the grey matter route of projections from the injection site to one of the projection targets. The direction of axonal flows is shown by cyan arrows in each panel judged by the angle of axon fibers and relative positions. The number on the top right indicates the section positions of the image in reference to the injection site (inj) in in mm (the number increases to the posterior direction). The areas of injection and targets was judged as in methods. The Allen data ID for these images are as follows (also see Table 1): (A) 126909424; (B) 141603190; (C) 114290938; (D) 126908007; ((E) 126861679; (F) 112791318; (G) 126907302; (H) 146858006; (I) 309004492; (J) 117298988; (K) 139426984; (L) 100148142. Bar: 250 μm.

**Fig. 4.**
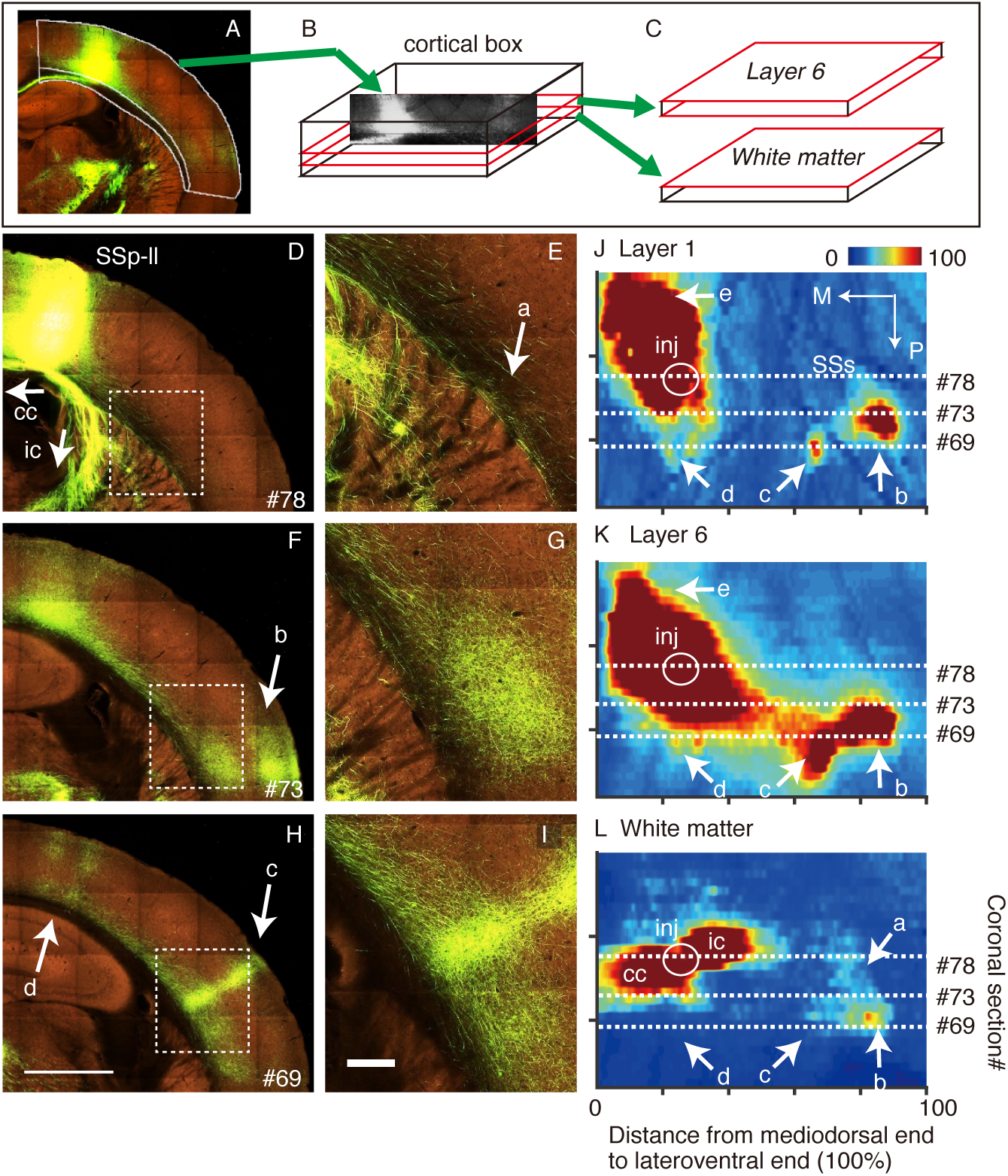
Cortical Box representation of the axonal spread for SSp-ll injection. (A-C) The schematic view of cortical box method. Example of a cortical section of SSp-ll (Allen data ID112791318; see Table 1) targeted image (A). The mediodorsal end, lateroventral end, inner contour, and outer contour were manually determined to select the part of the cortex for further processing. The selected cortical region was converted into a standard rectangle (B). The same procedure was repeated for a set of coronal sections to reconstruct 3D image of the cortex (cortical box). A specific layer fractions (e.g,. layer 6 and white matter) were extracted to demonstrate axonal spread in two-dimensional flatmaps (standardized layer maps). (D-L) Examples of the original coronal section images (D, F H) and the magnified cortical areas (E, G, I) indicated by dotted rectangles in the left panel were shown for comparison with the standardized layer maps for layer 1, 6 and white mater (J-L). The numbers in the right-bottom in D,F,H indicate the coronal section serial number from Allen Mouse Brain Connectivity atlas, of which corresponding positions were indicated in the standardized layer maps J-L. Injection site (inj) is determined by the extent of strong cell body labeling in the blue channel. The spots indicated by arrows b-d in flatmpas are columnar innervations that correspond to those in panels F and H. The weak signal in the white matter flatmap (panel L) indicated by arrow a include corticostriatal fiber shown in panel E. Scale bars for panels D, F, H; 1 mm, for panels E, G, I: 200µm. cc: corpus callosum. ic: internal capsule.

In the case of Fig. 3G, AAV injection into SSp-bfd led to axonal projection of both lateral (to AUDp) and medial (to RSPd) directions. Similar to Fig. 3A-F, lateral projections spread within the cortical grey matter to reach AUDp. To the medial side, there was a thick axon bundle in the white matter that continues to the contralateral side. However, this axon bundle did not branch to innervate area RSPd. Instead, many axon fibers spread parallel to the cortical surface. In particular, those in layer 6 continued to the bottom of area RSPd and turned upside to innervate across layers (Fig. 3G, right panel). Fig. 3H-J represent other examples of medial projections. Similar to Fig. 3G, medial projections proceeded parallel to the cortical surface within the grey matter to reach the target area. In all these cases, we observed axon fibers to run through layer 6 from the injection site to the target.

Fig. 3K and L represent two cases, in which AAV tracer was injected to the medial wall of the cortex. When AAV tracer was injected into ACA, we observed GFP signals in layer 6 to persist along AP axis (white arrow), until they finally spread out across layers to innervate RSPd and PTLp (Fig. 3K). This is another good example of axon fibers to pass through layer 6 to reach the distant target. When AAV tracer was injected into RSP, a thick axon bundle projected laterally within the white matter above the hippocampus. This axon bundle connects to the internal capsule in the more anterior sections. At this level, however, we were able to observe many axon fibers to branch off to enter the cortical grey matter lying above (white arrow). So, this is an exceptional example of axon fibers traveling through the white matter to reach the ipsilateral cortical areas. As far as we examined, RSP was the only area to take “white matter route” to reach the distant target. Once the axon fibers entered the grey matter, they spread widely within the cortical layers until they reach the final targets as in other examples.

Fig. 3 also illustrates variability of laminar patterns of cortical innervation. A frequently observed pattern is the columnar projections, such as from MOp to SS-s (Fig. 3A), SSp-m to SS-s (Fig. 3C), SSp-bfd to AUDp (Fig. 3G), or VISp/VISal to AUDd/PTLp (Fig. 3I). Quite often, innervation in layer 1 spread wider than the column size, suggesting horizontal spread within layer 1. In some cases, axon fibers spread widely across different areas within layer 1 with little or no innervation in layers 2–6 (e.g., Fig. 3B, Fig. 3H, Fig. 3K, Fig. 3L). On the other hand, such spread in layer 1 is not always observed (e.g., Fig. 3J). To accurately understand the laminar specificity, however, it is necessary to examine the spread of axon projections in 3D, and not by a single section. We come back to this topic later.

### Visualization of cortical connections by Cortical Box method

So far, we have described the patterns of cortical connectivity using “snapshots” out of a series of section image data. To follow trajectory of axon fibers across sections, we employed Cortical Box method that we previously devised to characterize laminar and area architecture of rat cortices (Hirokawa et al. 2008a; Hirokawa et al. 2008b; Watakabe 2012). In this method, a strip of dorsolateral cortical areas is first segmented out of the original image for transformation (Fig. 4A). At this point, the white and grey matter is also distinguished (see materials and method section for criteria of distinction). This strip is converted to a rectangle and the series of the rectangles are converted to a box. By re-slicing this box at particular lamina positions, we can have layer-specific flatmaps for the spread of axonal fluorescent signals (Fig. 4C).

Fig. 4 shows an example of Cortical Box representation of AAV tracer injection into SSp-ll (Allen data ID112791318; see Table 1). In the original coronal section images (Fig. 4D, F and H), the axonal projections originating from the injection site (Fig. 4D) were observed to proceed posterolaterally within layer 6 to reach two columnar foci (Fig. 4F and H, arrows b and c). We used 28 of such images (section #63–90) to reconstruct the Cortical Box and re-sliced them to present the flatmaps for layer 1, layer 6 and the white matter (Fig. 4J-L).

Now, the intense tracer signals in and around the injection site as well as the two conspicuous columnar foci described above were very well visualized in layer 1 flatmap (Fig. 4J; circle inj, arrows b and c). In addition, two weak neighboring columns shown in Fig. 4H (denoted by arrow d) were also visualized as two weak spots in this flatmap. For all these spots, axon signals were present at the corresponding location in layer 6 flatmap (see arrows b-d) as well, suggesting that axons projected vertically from layer 6 to layer 1 (Fig. 4K). One big difference between layer 1 and layer 6 flatmaps, however, was that tracer signals covered wider areas in layer 6. Actually, they were connected with each other to form pathways from the injection site to each projection target. In contrast, we did not observe such pathways between the injection site and spots b, c and d within the white matter (Fig. 4L).

In the white matter, very strong tracer signals were only present in positions that correspond to the axon fibers that proceed toward the corpus callosum (Fig. 4D, 4L; cc) and those toward the internal capsule (Fig. 4D, 4L; ic). We did observe some weak signals to continue from the internal capsule signals toward the lateral direction (see Fig. 4E and G). At least, a part of these signals come from the white matter fibers that enter the striatum (Fig. 4E, denoted by arrow a). Nevertheless, we cannot deny the possibility that a minor population of white matter axon fibers reached spots b and c through this pathway. Either way, it is clear that the majority of cortical projections proceed within the grey matter and very few, if any, axonal fibers use the white matter route to reach distant targets.

### Flatmap analyses demonstrate the predominance of “grey matter route” for cortical projections

In the present section, we examine the areal distribution patterns of tracer signals for 17 injections listed in Table 1 (also see Fig. 9E for topological localization), using layer-specific flatmaps as we did in the previous section. In Fig. 5, AAV tracer was injected in the medial sides, whereas in Fig. 6, AAV tracer was injected in the lateral sides. Fig. 7 presents four cases of medial wall injection, which were analyzed by a modified version of Cortical Box, namely, “Medial Box” method (see below for more details). Generally speaking, the patterns of layer 1 and layer 6 flatmaps were similar. In contrast, the robust tracer signals in layer 6 flatmap were very different from those of white matter flatmaps. Strong white matter signals corresponded to those that targeted corpus callosum and internal capsule, which ran parallel to the coronal sections (appears horizontal in the flatmap). Layer 6 signals often spread oblique to such horizontal lines and the overlay of the two flatmaps exhibited little overlap except around the injection site (bottom panels in Fig. 5–7).

**Fig. 5.**
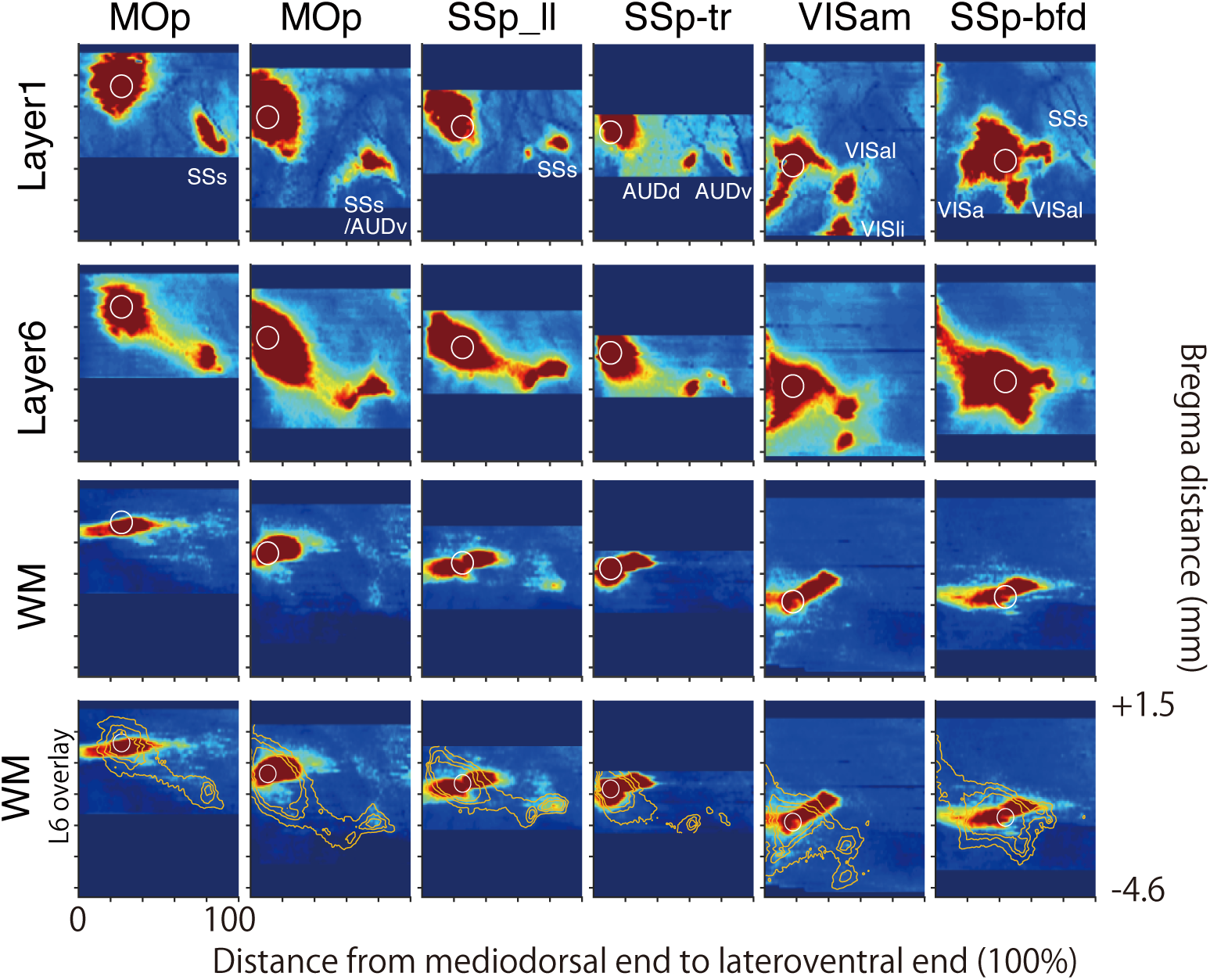
Spatial distribution of axonal spread in standardized cortical flatmaps (1) Standardized layer flatmaps for 6 different AAV tracer injections. Layer 1, 6 and white matter flatmaps were shown as typical examples. In order to compare different projections in the same coordinate, the anterior-posterior positions of each dataset were identified based on anatomical landmarks in reference to the Paxinos and Watson (2007) atlas (see Methods). Although we did not perform more precise registration, the Y position in each layer flatmap is considered to roughly correspond to the Bregma distance shown on the right. The dark blue space in the map indicates the area where we did not perform Cortical Box transformation. The white circles indicate the injection positions of AAV tracer. The bottom figures are the same as white matter map except that the contour of the signal from layer 6 map was overlaid as orange lines for comparison.

**Fig. 6.**
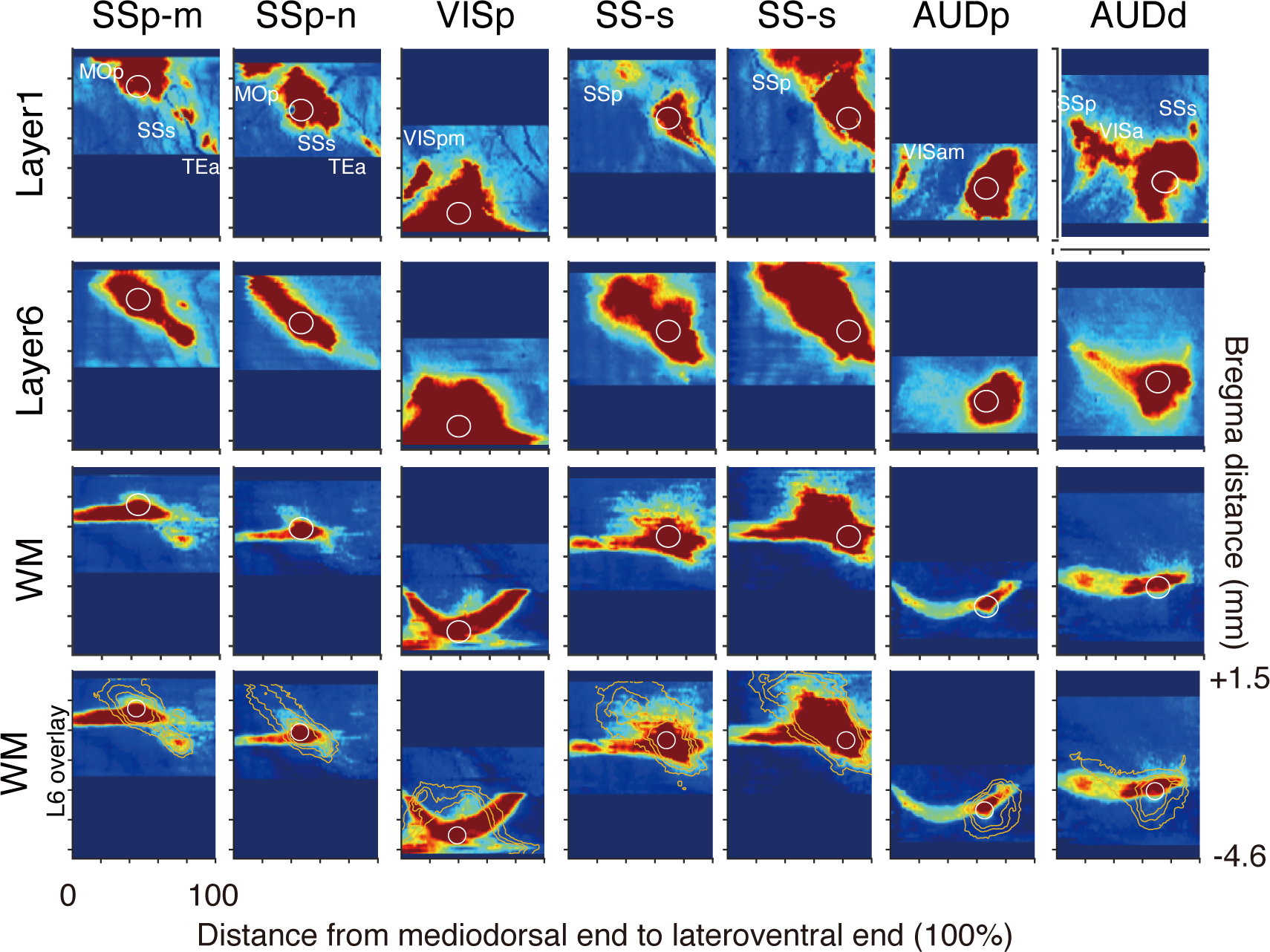
Spatial distribution of axonal innervation in standardized cortical flatmaps (2) Standardized layer flatmaps for 6 different AAV tracer injections. The conventions used in this figure are the same as those in with Figure 5.

**Fig. 7:**
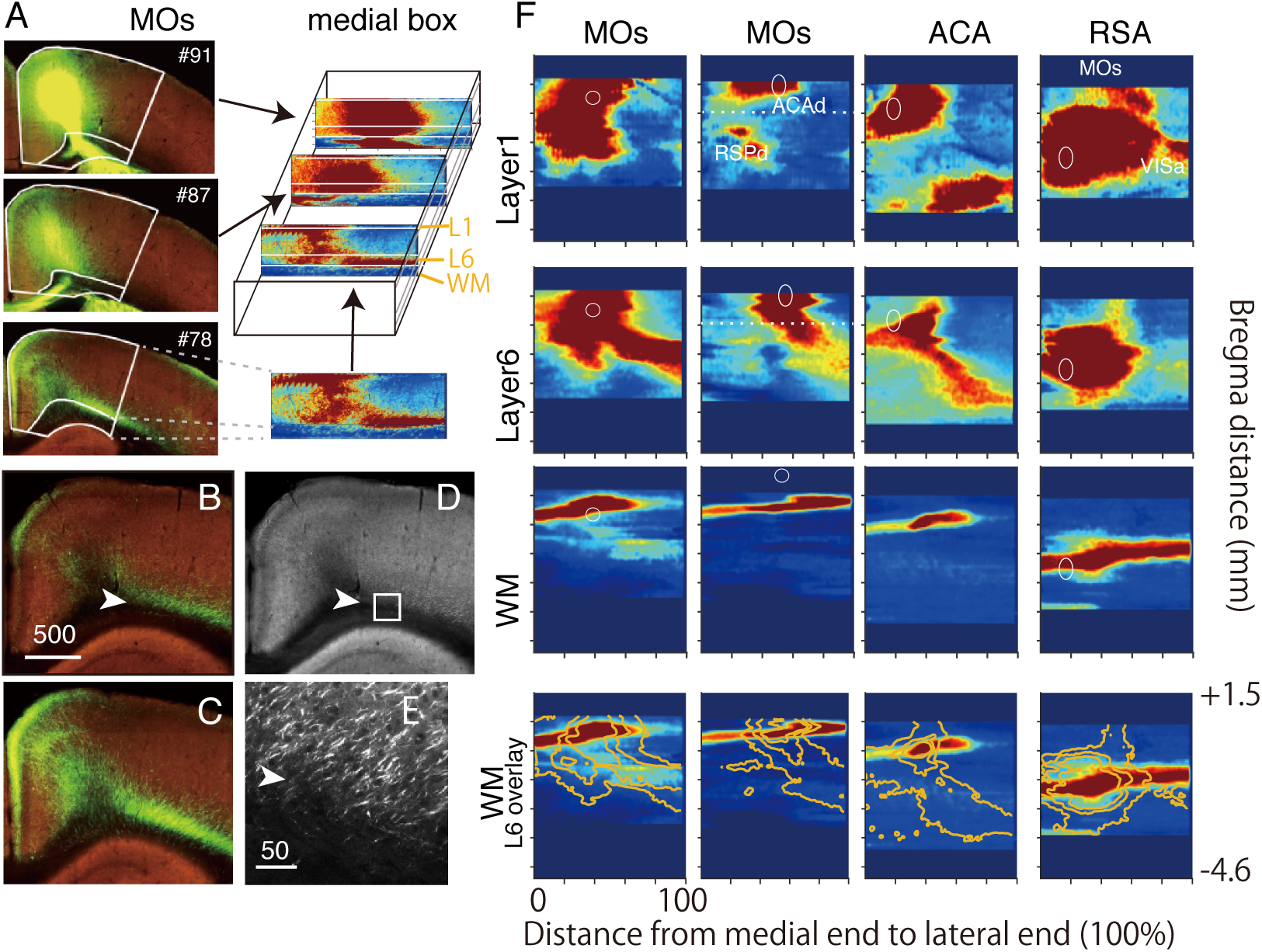
Analysis of axonal projections around the medial wall. (A) The schematic view of Medial Box method, which is modified from standard Cortical Box method. Example of cortical sections of injection into MOs (Allen data ID: 180916954). The medial end, lateral end, inner contour, and outer contour were manually determined to select the part of the cortex for further processing. The selected cortical region was converted into a standard rectangle. The same procedure was repeated for a set of coronal sections to reconstruct 3D image of the cortex. (B-E) Examples of original fluorescent images of MOs injections to show the absence of the white matter route for axonal spread to the posterolateral direction. Panels B and C show the images of similar AP position taken from Allen Data ID 141603190 and 180916954, respectively. Panel D shows the red channel view of panel B, which is magnified in panel E (square box). The border between layer 6 and the white matter (arrowhead) can be unambiguously identified by the presence of cell shadows in layer 6. Note that the intense axonal signals are clearly within layer 6. Scale Bars for panels B-D; 500 μm, for panel E; 50 μm. (F) Standardized layer flatmaps for six different AAV tracer injections. Same convention as in Figures 5 and 6.

Such difference was most conspicuous for six cases of medial injections shown in Fig. 5, but was similarly observed for seven cases of lateral injections in Fig. 6 in which stronger white matter signals were observed compared to Fig. 5. Because the white matter becomes thin and indistinct in the more lateral areas, a part of white matter signals may be contamination of layer 6 signals in these cases. In the injection of AUDp, we did not observe clear pathway of tracer signals that can account for the presence of an island of signals in layer 1 of VISam (Fig. 6). In this example, transduction of AAV was restricted to the lower layers and we were able to observe fine fibers to originate from the transduced cells to reach layer 1 of VISam in coronal section view (Fig. 8D). So, this is another example of axonal projection within the grey matter, although the axonal spread was not restricted to layer 6 and not very visible in the flatmap.

We also showed the “Medial Box” representation of four cases in Fig7. Because of higher level of distortion, the alignment became inascurate in these cases. However, the difference between layer 6 and white matter was still very clear. For example, in two cases of MOs injection, we observed posterolateral cortical paths within layer 6 that continued even out of the medial box (Fig. 7F). In the original images, the axon fibers were confirmed to be mostly restricted within the grey matter (Fig. 7B-D). The distinction between the grey and white matter was quite clear in this case, because we could identify shadows of neuronal cells only in the grey matter (Fig. 7E). We obtained similar results for both MOs injections, although the appearance of layer 1 flatmaps was different due to different levels of signal saturation (compare Fig. 7B and C). In the case of RSA injection, the lateral spread of tracers into VISa coincided with the extension of white matter fiber that targets the internal capsule. So, comparison of the flatmaps alone cannot determine the route of innervation. Regarding this point, we have already observed in Fig. 3L that axons toward VISa originated from the white matter. As far as we examined, this was the only clear case of white matter route innervation. To conclude, flatmap analyses demonstrated the predominance of “grey matter route” over “white matter route” for cortical projections.

### Front views of Cortical Box provide laminar information of axonal projections

While flatmaps provide the overview of the 2D spread of cortical projections, we can use the “front view” of the Cortical Box to visualize the laminar spread. In Fig. 8, we provide five example injections to show the front views in conjunction with the layer 1 flatmaps. The strength of the Cortical Box method is that we can combine the data distributed over several slices as maximum intensity projection (MIP). First, we show the identification of the injection centers. Because of high level of GFP expression, the green and red channels are saturated. But in the blue channel, we were able to find a threshold where we can identify individual cell bodies but not neurites coming out of the infected cells. MIP images of signals above such threshold provide the exact center of AAV transduction (Fig. 8, shown in blue in MIP panels). For SSp-m, MOp, and VISam injections (Fig. 8A, B and E), the transduced cells covered both upper and lower layers, whereas only the lower layers were infected in SSs and AUDp (Fig. 8C and D).

**Fig. 8.**
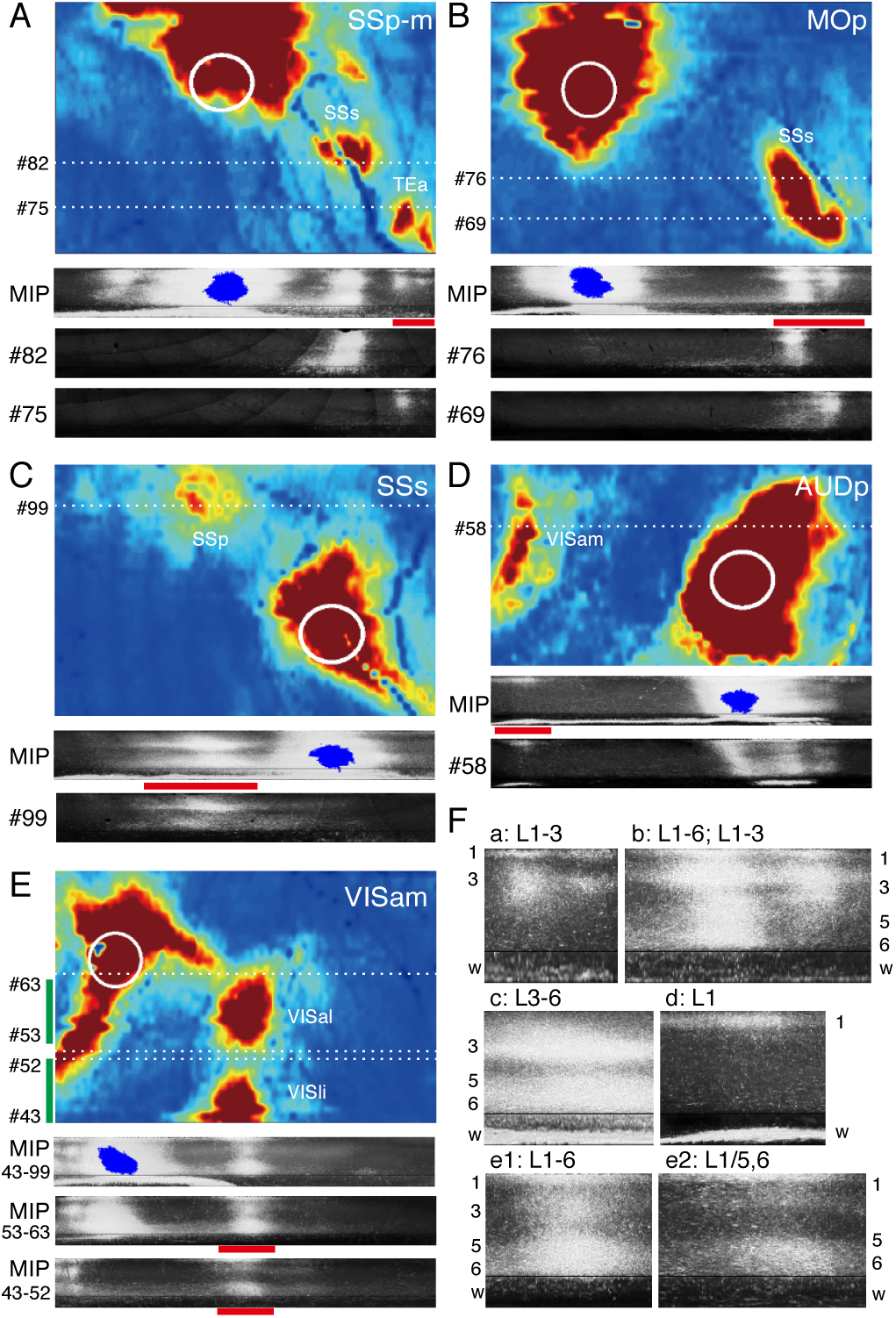
Front views of Cortical Box reveals layer-specific axonal projections. Maximum Intensity Projections (MIP) for the front view of the Cortical Box visualizes the axonal projections that encompass multiple sections. (A-E) Standardized flatmaps for layer 1 of SSp-m, MOp, SSs, AUDp and VISam injections are shown together with the MIP of front views. The standardized rectangle generated from coronal section images of the original Allen atlas are also indicated with the section numbers. The injection centers determined by the cell body signals in the blue channel are shown in blue for MIP front views. The red bar below the MIP views are magnified in panel F. For panel E, the MIPs for all the analyzed sections (43–99) are shown together with the MIPs for a part of the data (53–63, 43–52). (F) The MIP front views shown by red bar in panels A-E are magnified to show different laminar patterns of terminal extensions at remote areas.

Next, we examined the axonal pathways from the injection centers to the peripheral target areas. Although the analyzed data do not have the resolution to warrant continuity of axons at single fiber level, it is likely that the axons pass through regions with densest tracer signals. Generally speaking, the vicinity of the injection site was saturated with tracer signals that span all layers. These signals come from axon fibers and dendrites that extended only within the grey matter. In addition, tracer signals were identified in remote areas, which look like islands in the layer 1 flatmaps. The front view MIPs displayed how such “islands” are connected with the injection centers. A typical case was the connection through layer 6 pathway, such as between SSp-m and SSs (Fig. 8A; MIP) or between MOp and SSs (Fig. 8B; MIP). In these cases, we found the tracer signals to be continuous through layer 6 in the MIP view, although they are partially visible in each single section (e.g., compare section #76 and MIP of Fig. 8B). In VISam injection, we made two partial MIPs to visualize VISal and VISli projections separately (Fig. 8E). Interestingly, we observed dense tracer signals in layer 6 for both MIP views, suggesting the existence of axon fibers that spread widely within layer 6. In AUDp injection (Fig. 8D), we could not observe clear axonal pathway in the front view MIP, despite presence of dense terminations in layer 1 of VISam. Similarly, we did not observe clear axonal pathways for SSp-m injection between SSs and TEa (Fig. 8A). In such cases, thin axon fibers were found dispersed in the cortical grey matter in the original images. These examples suggest that area-to-area connections can be conveyed through dispersed axons within the grey matter.

Finally, we examined the laminar specificity of axon terminal convergence in the remote areas (Fig. 8F). In some cases, we observed concentration of tracer signals in layer 6, which is continuous from the “pathway signals” and accompanied concentration of tracer signals in the upper layers as well (Fig. 8F; b, c, e1 and e2). In other examples, we observed concentration of tracer signals in the upper layers with only passing fibers in layers 5 and 6 (Fig. 8F, a, b and d). Innervation in layer 1 was almost always observed, but in one case, we observed no innervation in layer 1 with concentrated tracer signals in layers 3/4 and 5/6 (Fig. 8F; c). These examples illustrate the variability of axonal terminations in various cortical projection patterns. The front view MIPs for all the 17 Cortical Boxes that we analyzed are shown in Supplementary Fig.1.

### Quantitative analyses of layer-specific cortical projection patterns

To more systematically analyze the layer specificity of intracortical axonal projections, we decomposed the Cortical Box into columnar modules and investigated the laminar distribution of tracer signals in each module. By this procedure, the 3D distribution of tracer signals within the Cortical Box was translated to the 2D distribution of columnar modules, each of which is characterized by a unique laminar profile. To objectively classify these laminar profiles, we conducted clustering analysis on the columnar modules from 17 Cortical Boxes. We first estimated how many different laminar patterns are appropriate to classify the tracer distributions. To this end, we systematically changed the number of clusters used for the clustering analysis and measured the similarities among the resulting clusters (see Methods). In this analysis, we found that the total intra-cluster variation reaches the first plateau at five clusters, suggesting that further increase in cluster number does not improve the clustering (Fig. 9A). As Fig. 9B shows, the resulting 5 clusters exhibited unique laminar patterns. Among them, the pattern “e” represents all-layer pattern, which is often found in the vicinity of injection centers. Patterns “a-d” exhibited more or less layer specific patterns (Fig. 9B). The actual example columns that were classified into these clusters are shown in Fig. 9C. To test the robustness of this classification, we performed the clustering using only a subset of the 17 injections and compared the obtained patterns with the original pattern. As shown in Supplementary Fig.3, significant similarity to the original (N=17) patterns were observed only by using two samples (N=2) for clustering and the correlation coefficient reached a high value (>0.9) when six datasets were used. This result suggests that the five patterns shown in Fig. 9B is quite robust, and that comparable layer patterns will still be obtained, independent of dataset selection (Supplementary Fig.3).

**Fig. 9.**
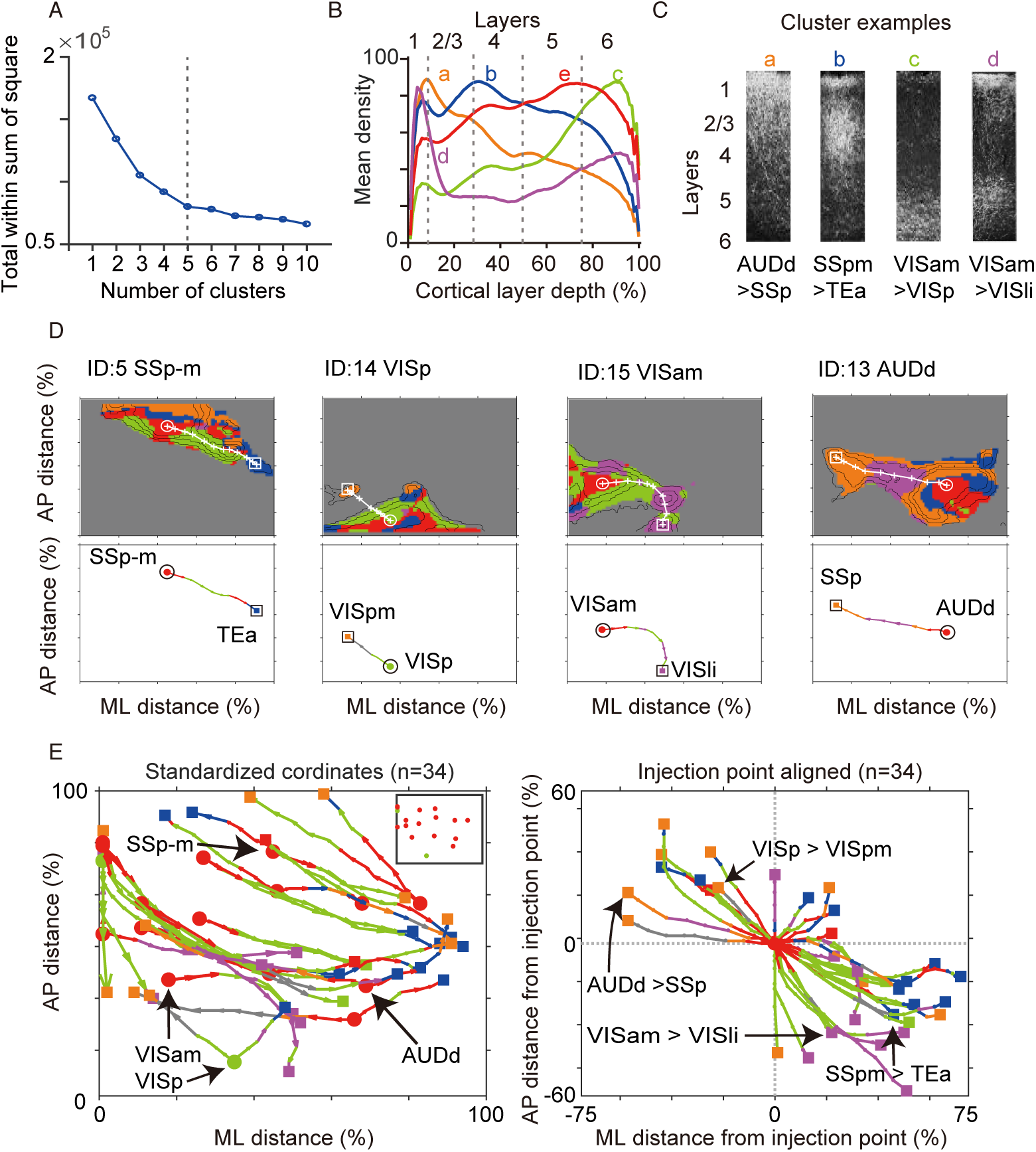
lustering analysis of layer distributions. (A) Clustering analysis for all columns of cortical box from 17 injection samples. Similarities of clusters plotted as a function of cluster numbers used for the clustering analysis (see Methods). (B) The averaged layer distributions of 5 clusters (patterns “a-e”) from the pial surface (0% of cortical depth) to the cortex/white matter border (100%). (C) Examples of actual layer pattern for each cluster. (D) Examples of cluster assignment in horizontal layer map. The colors of the areas correspond to the cluster identity shown in (B). A white circle and a rectangle in each map represent the tracer injection site and the furthest projection areas. The projection pathways connecting those two areas were indicated by white crosses and lines. The cluster identities on each projection pathway were indicated by line colors in the bottom. (E) The projection pathways (n=34) from 17 injection samples (2 pathways for each sample) were overlaid in a map (left panel). The inset indicates the 17 injection sites. The each projection pathway was aligned so that the injection point comes to the center of the figure (right panel).

To visualize the spatial distributions of thus-classified columnar modules, we overlaid the color-coded identities on the flatmaps (Fig. 9D, Supplementary Fig.2). In Fig. 9D, we showed four examples of such color-coded “laminar maps”. These laminar maps provided rich information on how axonal signals spread from the injection site to the surrounding areas.

First, as we mentioned above, pattern “e” (red) was often found near the injection site. Although the quantitation is compromised in the vicinity of injection site due to saturation of the tracer signals, we could clearly observe axons to extend in all layers to densely innervate the nearby areas. Second, the injection centers were often surrounded by pattern “c” (green), which is characterized by higher signals in layer 6, which is consistent with the front view analyses in Fig. 8. Third, in peripheral areas, patterns “a” (orange) and “d” (purple) were often found, which both exhibit enriched signals in layer 1, suggesting the spread of axons in layer 1. To more systematically compare the tracer spread for different samples, we paid attention to the projection pathways between the injection site and select target areas. The lower panels of Fig. 9D demonstrate such pathway analyses for the furthest target area in each sample. For example, in the case of SSp-m injection (Fig. 9D, ID 5), the injection site (SSp-m) showed pattern “e” (red circle) and the remote target area (TEa) showed pattern “b” (blue square). The most likely pathway for this projection (see methods section for tracking method) comprised mostly of pattern “c” (green lines). This suggests that the axons that originated from SSp-m mainly proceeded within layer 6 to reach TEa, where the innervation spread toward the upper layers. We selected two furthest target areas for each sample for this pathway analyses, and remapped on a single common flatmap for comparison.

As shown in Fig. 9E (left panel), the composite flatmap showed clear bias of projection profiles, despite random scattering of the injection sites (Fig. 9E, inset in left panel). First, we found that most of the long-range projections run along the anteromedial to posterolateral axis. This bias was captured more clearly by replotting the projections so that the injection site comes at the center of the plot (Fig. 9E, right panel, P=0.001, Kuiper test): whereas 14 and 13 projections were found within the anteromedial and posterolateral quadrants, respectively, only 7 projections were found within the anterolateral quadrant. No projection was found in posteromedial quadrant. Second, the laminar profiles of the long-range projections were dominated by pattern “c” (green), which means that the axons proceeded through layer 6. This feature was again better visualized by centering the injection site. Third, the laminar profiles at the projection terminals exhibited bias depending on the direction of projections; pattern “a” was more prevalent in the anteromedial quadrant, whereas patterns “b” and “d” were more prevalent in the posterocaudal quadrant.

In previous studies, Burkhalter and coworkers proposed that the axonal targeting of layers 2–4, and of layer 1 represent feedforward and feedback projections, respectively, based on the anterograde tracing of the rodent visual areas (Coogan and Burkhalter 1993; D’Souza et al. 2016·; D’Souza and Burkhalter 2017). We were interested in whether the same concept can be applied beyond the visual areas. In our clustering scheme, pattern “d” represented enrichment in layer1 and layer 6, pattern “a” represented enrichment in upper layers with the peak in layer 1 and pattern “b” represented enrichment in layers 2–4 with some contribution from layer 1. Therefore, the “a” and “d” patterns are considered to be “feedback”, whereas “b” pattern is considered to be “feedforward”, according to this scheme. As shown in Table 2, the projections originating from the primary areas tended to terminate with pattern “b” (12/18; 66.7%), whereas the projections originating from the higher order areas tended to terminate with patterns “a” or “d” (14/16; 87.5 %). The statistical significance of this difference was confirmed by bootstrap method (p<0.00005; see Methods). We also examined the occurrence probabilities of the four lamina patterns (except pattern “e”) for the entire cortex in each injection and compared between the primary areas and higher order areas (Supplementary Fig.3A). In both categories, pattern “c” was high, which is considered to be axon pathway. We observed statistical difference in patterns “b” and “d”, which were more abundant in the primary (p<0.05) and higher order areas (p<0.005), respectively. The differential occurrence probability of pattern “d”, which represents enrichment in layers 1 and 6, was especially striking. This is interesting, because the “feedback” projection of macaque visual cortices also targets layers 1 and 6 (Rockland and Pandya 1979; Felleman and Van Essen 1991). Thus, our data appear to suggest the existence of similar rule for area-specific innervation patterns between mouse and primates.

## IV. DISCUSSION

The main finding of the current study is the predominance of the “grey matter route” over the “white matter route” for ipsilateral cortico-cortical connectivity in the mouse brain. To our knowledge, this point has never been investigated in a systematic way. In our study, we investigated the projection patterns of 17 different injections, and successfully visualized the above finding by Cortical Box method. Furthermore, detailed laminar analyses revealed a complex architecture of the mouse projections within the grey matter. In the vicinity of the injection sites, we observed horizontal extension of the axon fibers in both upper and lower layers, which is similar to the primate intrinsic connections (Levitt et al. 1993; Lund et al. 1993). On the other hand, we also found point-to-point connectivity for long-range connections, which appeared as an island of intense tracer signals in the upper layer flatmap. The majority of such “islands” were connected via axons in layer 6, which is reminiscent of the primate’s extrinsic connections traveling through the white matter. Interestingly, the projections from the primary and higher-order areas to the distant targets preferentially terminated in layers 2–4 and layer 1, respectively, suggesting possible hierarchical organization similar to that of macaques (Rockland and Pandya 1979; Felleman and Van Essen 1991; Coogan and Burkhalter 1993; D’Souza et al. 2016; D’Souza and Burkhalter 2017). Based on these observations, we suggest that the cortico-cortical connectivity in the mouse brain also consists of “extrinsic” and “intrinsic”-type connections in a similar manner to primate connections, but that these two types are intermixed within the grey matter.

### Technical considerations

Our study showed that Allen Mouse Brain Connectivity atlas provides excellent data for the investigation of the mouse connectivity. We found, however, several shortcomings that required cautions. First, we found it difficult to examine fine morphological features of the axonal arbors with the current XYZ resolution. Although we were interested in whether the layer 6 axon fibers form synapses on the way to the remote target, the image data in the database was insufficient. To clarify this point, we performed an additional tracing and confocal imaging experiment to supplement the Allen data, and found axon fibers with and without boutons to coexist in layer 6. Second, the tracer signals of the database tended to be weaker in the white matter region, probably due to optical interference of highly myelinated tissue. Regarding this point, we believe that the effect was minimal, because we could detect even thin corticostriatal fibers within the white matter. Third, we noticed that many of the lateral AAV injections were restricted to the lower layers (e.g., see Fig. 8). This is probably because of limited dispersion of AAV by the iontophoretic injection (Oh et al. 2014). We did not pursue this problem further, because we could not find appropriate datasets to estimate this influence. More detailed studies using layer-specific transgenic lines may, in the future, solve this problem.

Cortical Box method was previously developed to standardize and quantify histological serial section data (Hirokawa et al. 2008a; Hirokawa et al. 2008b; Watakabe 2012). With a modification to distinguish the white and grey matter compartment, it proved to be a powerful tool to visualize and analyze the cortical axonal projections. The clustering analysis, in particular, provided an objective means to investigate the layer-specific axonal extensions. One pitfall of the current analysis is that the tracer signals are often saturated near the injection site and at some terminal regions. When the tracer signals are saturated, it affects the relative abundance even after normalization (e.g., see Fig. 7). This is a difficult problem to overcome, because it requires the improvement in various steps of labeling and imaging. Nevertheless, we believe that the effect of saturation is kept minimal thanks to systematic experimental design of the Allen dataset.

### Spatial organization of inter-areal connections in the mouse cortex

In Fig. 9E and F, we have shown the bias of cortico-cortical projections for long-range connections: the majority of these projections was aligned in the anteromedial-posterolateral (AM/PL) direction. What does this bias stand for? We can account for this bias from developmental and functional points of view. It is now well established that the regionalization of the mouse cortex occurs under the influence of gradients of molecules (Sansom and Livesey 2009). Interestingly, the direction of Pax6/Emx2 gradient is orthogonal to the current AM/PL direction. There is thus a possibility that axonal projections occur during development in reference to such gradients. Alternatively, the bias of axons may be a secondary effect of biased positioning of the primary and higher order areas, which are generated according to the molecular gradients.

From the functional point of view, our data suggests the existence of a “layer 6 cortical route” that flows in AM/PL direction. If this idea is correct, we would expect to see preferential coupling of activities in pairs of areas that are positioned in AM/PL direction. Regarding this point, GCaMP6f-expressing transgenic mice has been used to investigate which cortical regions exhibit temporally correlated activities in resting state (Vanni et al. 2017). In higher frequency mode (2–5 Hz), two main clusters of highly correlated activities were found: (1) a lateral/anterior cluster that encompasses all the somatosensorimotor areas and (2) a medial/posterior cluster that encompasses visual cortex and the retrosplenial cortex. This finding suggests the coupling of activity within each cluster, but not across the cluster, which topology matches well with our connectional map. Similar clusters have been found in resting state fMRI imaging (Liska et al. 2015) or in connectivity mapping using viral tracers (Zhang et al. 2016). At low frequency mode (0.1–1 Hz), slightly different functional clusters have been observed (Vanni et al. 2017), which may be dependent on slow-conducting fibers. Our structural analyses thus appear to be largely in agreement with the functional connectivity mapping.

One characteristic feature of the mouse cortical projections is that the mouse projections extend beyond modalities, which has been noted previously for V1 (Wang et al. 2012) and for other areas in this study (e.g., see Table 2). In contrast, the monkey areas generally do not cross modalities in the early areas (Felleman and Van Essen 1991; Goodale and Milner 1992; Markov et al. 2014 but see (Rockland and Ojima 2003; Ghazanfar and Schroeder 2006; VanAtteveldt et al. 2014). This difference is likely to be attributed to different brain size and total neuron numbers. It is suggested that cortical wiring is under the constraints to keep the amount of the connecting axon fibers to be minimum while maintaining functionality of the network (Kaas 2000; Wang et al. 2008; Bullmore and Sporns 2012; Hofman 2014). We suggest that the predominance of the grey matter route for inter-area connections as well as multimodal connectivity may be an adaptive strategy for a small-brained mammal like mice. The long-range connections within the grey matter in mice may provide additional opportunities to make synaptic connections on the way and maximize the use of smaller number of neurons.

### Laminar organization of inter-areal connections in the mouse cortex

The concept of hierarchical organization of the cortical areas was originally formulated based on segregated laminar targeting of visual area connectivity in the macaque monkey and was later extended to the entire cortical areas (Rockland and Pandya 1979; Felleman and Van Essen 1991). This influential concept was, nevertheless, not easily transferable to the rodent cortex. By detailed tracer experiments, Burkhalter and coworkers showed the existence of multiple visual areas receiving retinotopic inputs from V1 in rodents (Coogan and Burkhalter 1993; Wang and Burkhalter 2007; Wang et al. 2012), but the input layer specificity was somewhat different between rodents and primates (reviewed in D’Souza and Burkhalter 2017). They suggested that the ratio of layer 2–4 and layer 1 inputs as the conserved metric of cortical hierarchy and used it to demonstrate the hierarchical order of the visual areas (Coogan and Burkhalter 1993; D’Souza et al. 2016; D’Souza and Burkhalter 2017). With similar criteria for analysis, we examined datasets that cover the entire dorsal cortical areas and now found evidence for the existence of cortical hierarchy even outside the visual areas.

The present study, together with past studies (reviewed in Markov et al. 2014; D’Souza and Burkhalter 2017), strongly suggests the different roles of inputs into layers 2–4 and layer 1 in both primates and mice. One classic view of the role of layer-specific inputs is that those to layers 3–4 are “driving”, and those to layers 1 and 6 are “modulatory” (Crick and Koch 1998; Jones 1998; Sherman and Guillery 1998; DePasquale 2011). Recent mouse studies, however, suggest that the “top-down” inputs to layer 1 and deep layers have profound effects on somatosensory (Manita et al. 2015) and visual (Leinweber et al. 2017) sensations beyond simple modulatory effects. It is not clear at present how we can generalize these findings to other cortical connections or to connections in primates. Nevertheless, we believe that the mouse will continue to be a good model animal to study the cortical connectivity as long as we accurately understand the similarities and differences as were revealed in this study.

## ACKNOWLEDGMENT

This work was supported by JSPS KAKENHI Grant Number 16K07036 (to A.W.) and 16K18380 (to J.H.).

## REFERENCES

Bullmore E, Sporns O. 2012. The economy of brain network organization. Nat Rev Neurosci. 13: 336–49.

Coogan TA, Burkhalter A. 1993. Hierarchical organization of areas in rat visual cortex. J Neurosci. 13: 3749–772.

Crick F, Koch C. 1998. Constraints on cortical and thalamic projections: the no-strong-loops hypothesis. Nature. 391: 245–50.

De Pasquale R, Sherman SM. 2011. Synaptic properties of corticocortical connections between the primary and secondary visual cortical areas in the mouse. J Neurosci. 31: 16494–16506.

D’Souza RD, Burkhalter A. 2017. A Laminar Organization for Selective Cortico-Cortical Communication. Front Neuroanat. 11: 71.

D’Souza RD, Meier AM, Bista P, Wang Q, Burkhalter A. 2016. Recruitment of inhibition and excitation across mouse visual cortex depends on the hierarchy of interconnecting areas. Elife. 5

Felleman DJ, Van Essen DC. 1991. Distributed hierarchical processing in the primate cerebral cortex. Cereb Cortex. 1: 1–7.

Finlay BL. 2016. Principles of Network Architecture Emerging from Comparisons of the Cerebral Cortex in Large and Small Brains. PLoS Biol. 14:e1002556.

Ghazanfar AA, Schroeder CE. 2006. Is neocortex essentially multisensory? Trends Cogn Sci. 10: 278–85.

Goodale MA, Milner AD. 1992. Separate visual pathways for perception and action. Trends Neurosci. 15: 20–5.

Goulas A, Uylings HBM, Hilgetag CC. 2017. Principles of ipsilateral and contralateral cortico-cortical connectivity in the mouse. Brain Struct Funct. 222: 1281–295.

Hirokawa J, Bosch M, Sakata S, Sakurai Y, Yamamori T. 2008a. Functional role of the secondary visual cortex in multisensory facilitation in rats. Neuroscience. 153: 1402–417.

Hirokawa J, Watakabe A, Ohsawa S, Yamamori T. 2008b. Analysis of area-specific expression patterns of RORbeta, ER81 and Nurr1 mRNAs in rat neocortex by double in situ hybridization and cortical box method. PLoS One. 3:e3266.

Hofman MA. 2014. Evolution of the human brain: when bigger is better. Front Neuroanat. 8: 15.

Horvá S, GăăuţR, Ercsey-Ravasz M, Magrou L, GăăuţR, Van Essen DC, Burkhalter A, Knoblauch K, Toroczkai Z, Kennedy H. 2016. Spatial Embedding and Wiring Cost Constrain the Functional Layout of the Cortical Network of Rodents and Primates. PLoS Biol. 14:e1002512.

Jones EG. 1998. A new view of specific and nonspecific thalamocortical connections. Adv Neurol. 77:49–1; discussion 72–3.

Kaas JH. 2000. Why is brain size so important: Design problems and solutions as neocortex gets biggeror smaller. Brain and Mind. 1: 7–3.

Leinweber M, Ward DR, Sobczak JM, Attinger A, Keller GB. 2017. A Sensorimotor Circuit in Mouse Cortex for Visual Flow Predictions. Neuron. 95:1420–432.e5.

Levitt JB, Lewis DA, Yoshioka T, Lund JS. 1993. Topography of pyramidal neuron intrinsic connections in macaque monkey prefrontal cortex (areas 9 and 46). J Comp Neurol. 338: 360–76.

Liska A, Galbusera A, Schwarz AJ, Gozzi A. 2015. Functional connectivity hubs of the mouse brain. Neuroimage. 115: 281–91.

Lund JS, Yoshioka T, Levitt JB. 1993. Comparison of intrinsic connectivity in different areas of macaque monkey cerebral cortex. Cereb Cortex. 3: 148–62.

Manita S, Suzuki T, Homma C, Matsumoto T, Odagawa M, Yamada K, Ota K, Matsubara C, Inutsuka A, Sato M, Ohkura M, Yamanaka A, Yanagawa Y, Nakai J, Hayashi Y, Larkum ME, Murayama M. 2015. A Top-Down Cortical Circuit for Accurate Sensory Perception. Neuron. 86: 1304–316.

Markov NT, Vezoli J, Chameau P, Falchier A, Quilodran R, Huissoud C, Lamy C, Misery P, Giroud P, Ullman S, Barone P, Dehay C, Knoblauch K, Kennedy H. 2014. Anatomy of hierarchy: feedforward and feedback pathways in macaque visual cortex. J Comp Neurol. 522: 225–59.

Oh SW, Harris JA, Ng L, Winslow B, Cain N, Mihalas S, Wang Q, Lau C, Kuan L, Henry AM, Mortrud MT, Ouellette B, Nguyen TN, Sorensen SA, Slaughterbeck CR, Wakeman W, Li Y, Feng D, Ho A, Nicholas E, Hirokawa KE, Bohn P, Joines KM, Peng H, Hawrylycz MJ, Phillips JW, Hohmann JG, Wohnoutka P, Gerfen CR, Koch C, Bernard A, Dang C, Jones AR, Zeng H. 2014. A mesoscale connectome of the mouse brain. Nature. 508: 207–14.

2014. The rat nervous system, fourth edition.

Paxinos G, Franklin KBJ. 2004. The mouse brain in stereotaxic coordinates. Elsevier Academic Press: Amsterdam; Boston.

Paxinos 2014. The rat nervous system, fourth edition. London (UK): Academic Press

Ragan T, Kadiri LR, Venkataraju KU, Bahlmann K, Sutin J, Taranda J, Arganda-Carreras I, Kim Y, Seung HS, Osten P. 2012. Serial two-photon tomography for automated ex vivo mouse brain imaging. Nat Methods. 9: 255–58.

Rockland KS, Ojima H. 2003. Multisensory convergence in calcarine visual areas in macaque monkey. Int J Psychophysiol. 50: 19–6.

Rockland KS, Pandya DN. 1979. Laminar origins and terminations of cortical connections of the occipital lobe in the rhesus monkey. Brain Res. 179: 3–0.

Sansom SN, Livesey FJ. 2009. Gradients in the brain: the control of the development of form and function in the cerebral cortex. Cold Spring Harb Perspect Biol. 1:a002519.

Schü A, Chaimow D, Liewald D, Dortenman M. 2006. Quantitative aspects of corticocortical connections: a tracer study in the mouse. Cereb Cortex. 16: 1474–486.

Sherman SM, Guillery RW. 1998. On the actions that one nerve cell can have on another: distinguishing “drivers” from “modulators”. Proc Natl Acad Sci U S A. 95: 7121–126.

Tamamaki N, Nakamura K, Furuta T, Asamoto K, Kaneko T. 2000. Neurons in Golgi-stain-like images revealed by GFP-adenovirus infection in vivo. Neurosci Res. 38: 231–236.

van Atteveldt N, Murray MM, Thut G, Schroeder CE. 2014. Multisensory integration: flexible use of general operations. Neuron. 81: 1240–1253.

Vanni MP, Chan AW, Balbi M, Silasi G, Murphy TH. 2017. Mesoscale Mapping of Mouse Cortex Reveals Frequency-Dependent Cycling between Distinct Macroscale Functional Modules. J Neurosci. 37: 7513–533.

Wang Q, Burkhalter A. 2007. Area map of mouse visual cortex. J Comp Neurol. 502: 339–57.

Wang Q, Sporns O, Burkhalter A. 2012. Network analysis of corticocortical connections reveals ventral and dorsal processing streams in mouse visual cortex. J Neurosci. 32: 4386–399.

Wang SS-H, Shultz JR, Burish MJ, Harrison KH, Hof PR, Towns LC, Wagers MW, Wyatt KD. 2008. Functional trade-offs in white matter axonal scaling. J Neurosci. 28: 4047–056.

Watakabe A, Hirokawa J, Ichinohe N, Ohsawa S, Kaneko T, Rockland KS, Yamamori T. 2012. Area-specific substratification of deep layer neurons in the rat cortex. J Comp Neurol. 520: 3553–3573.

Watakabe A, Takaji M, Kato S, Kobayashi K, Mizukami H, Ozawa K, Ohsawa S, Matsui R, Watanabe D, Yamamori T. 2014. Simultaneous visualization of extrinsic and intrinsic axon collaterals in Golgi-like detail for mouse corticothalamic and corticocortical cells: a double viral infection method. Front Neural Circuits. 8: 110.

Zhang S, Xu M, Chang W-C, Ma C, Hoang Do JP, Jeong D, Lei T, Fan JL, Dan Y. 2016. Organization of long-range inputs and outputs of frontal cortex for top-down control. Nat Neurosci. 19: 1733–742.

Zhang ZW, Deschêes M. 1997. Intracortical axonal projections of lamina VI cells of the primary somatosensory cortex in the rat: a single-cell labeling study. J Neurosci. 17: 6365–379.

Zingg B, Hintiryan H, Gou L, Song MY, Bay M, Bienkowski MS, Foster NN, Yamashita S, Bowman I, Toga AW, Dong H-W. 2014. Neural networks of the mouse neocortex. Cell. 156: 1096–111.

